# Linking *in vivo* muscle dynamics to *in situ* force-length and force-velocity reveals that guinea fowl lateral gastrocnemius operates at shorter than optimal lengths

**DOI:** 10.1101/2023.10.11.561922

**Authors:** MJ Schwaner, DL Mayfield, E Azizi, MA Daley

## Abstract

Force-length (F-L) and force-velocity (F-V) properties characterize skeletal muscle’s intrinsic properties under controlled conditions, and it is thought that these properties can inform and predict *in vivo* muscle function. Here, we map dynamic *in vivo* operating range and mechanical function during walking and running, to the measured *in situ* F-L and F-V characteristics of guinea fowl (*Numida meleagris*) lateral gastrocnemius (LG), a primary ankle extensor. We use *in vivo* patterns of muscle tendon force, fascicle length, and activation to test the hypothesis that muscle fascicles operate at optimal lengths and velocities to maximize force or power production during walking and running. Our findings only partly support our hypothesis: *in vivo* LG velocities are consistent with optimizing power during work production, and economy of force at higher loads. However, LG does not operate at lengths on the force plateau (±5% Fmax) during force production. LG length was near L_0_ at the time of EMG onset but shortened rapidly such that force development during stance occurred almost entirely on the ascending limb of the F-L curve, at shorter than optimal lengths. These data suggest that muscle fascicles shorten across optimal lengths in late swing, to optimize the potential for rapid force development near the swing-stance transition. This may provide resistance against unexpected perturbations that require rapid force development at foot contact. We also found evidence of passive force rise (in absence of EMG activity) in late swing, at lengths where passive force is zero *in situ*, suggesting that dynamic history dependent and viscoelastic effects may contribute to *in vivo* force development. Direct comparison of *in vivo* work loops and physiological operating ranges to traditional measures of F-L and F-V properties suggests the need for new approaches to characterize dynamic muscle properties in controlled conditions that more closely resemble *in vivo* dynamics.

## INTRODUCTION

The isometric force-length (F-L) and isotonic force-velocity (F-V) relationships are two widely used empirical measures of muscle contractile properties that characterize mechanical output under controlled conditions. The most common purpose of isometric F-L and isotonic F-V experiments is to systematically characterize muscle fascicle contractile properties under consistent conditions that can be compared across fiber types, muscles, species, and experimental treatment groups (e.g., changes with training, unloading, age, drug treatment, etc.) (Always 1995; Rassier et al., 1999; Jones 2010; Raj et al., 2010). The isometric F-L and isotonic F-V relationships demonstrate optimal operating ranges for muscle force and power production (e.g., Fenn and Marsh, 1935; Gordon et al., 1966; James et al., 1998; Zajac, 1989; Askew and Marsh, 1998), which are used to interpret *in vivo* muscle fascicle lengths, velocities, and forces. These properties are also widely used to specify muscle actuator properties in musculoskeletal and neuromechanical simulations of movement, in software such as OpenSim, AnyBody and others (e.g., Delp et al. 2007; Biewener et al., 2014; Seth et al., 2018; Cox et al., 2019). Considering the wide-spread use of the F-L and F-V properties for characterizing muscle fascicle contractile performance, it is important to understand how these properties relate to muscle fascicle operating ranges and force and power output *in vivo*.

The isometric F-L relationship finds its origin in the requirement of the muscle’s contractile proteins, actin, and myosin, to overlap to enable cross-bridge attachment and force production (Blix 1894; Gordon et al., 1966; Huxley 1957). The isometric F-L relationship is created through a series of fixed end tetanic isometric contractions over a series of lengths (Blix 1894; Gordon et al., 1966; Huxley 1957). In this relationship, force is optimal at intermediate lengths where overlap between actin and myosin can form the maximum number of cross-bridges. Muscles can operate *in vivo* over a wide range of lengths (or strain) spanning different regions along the isometric F-L curve. There is no consensus on typical operating lengths as previous studies have shown muscles operating on the ascending the plateau or the descending limb (e.g., Lieber and Boakes 1988; Herzog et al., 1992; Burkholder and Lieber 2001; Maganaris 2001; Ahn et al., 2006; Rubenson et al. 2012; Gidmark et al., 2013; Azizi 2014, Ahn et al., 2018). Relating the isometric F-L properties to *in vivo* function is made more difficult by the fact that few studies measuring *in vivo* force map those patterns to the F-L curve in the same muscle. Some previous studies have compared isometric F-L relationship to *in vivo* operating ranges, but most of these studies focused on *in vivo* strain ranges, without direct measures of muscle force (e.g., Gregor et al., 1988; Holt and Azizi, 2014; Ahn et al., 2018). The static nature of the F-L curve implies that this relationship is not applicable to the dynamic conditions of *in vivo* muscle measures; nonetheless, the relationship continues to be widely used in “Hill-type” model simulations of musculoskeletal dynamics. It remains unknown whether muscles operate on the force plateau of the isometric F-L curve during peak force development of in vivo dynamic, submaximal contractions.

The binding and detachment of cross-bridges in a cyclic pattern result in the isotonic F-V relationship, in which shortening velocity decreases with increasing steady-state isotonic load, resulting in an inverse hyperbolic relationship between force and shortening velocity (Hill 1938; Caiozzo 2002; Nelson et al., 2004; Sugi and Ohno 2019). Maximal force capacity occurs during isometric (zero velocity) contractions (Hill 1938: Alcazar et al., 2019). The isotonic F-V relationship is obtained by completing a series of isotonic contractions and is considered a foundational characteristic of skeletal muscle physiology. It was historically developed to test theories of cross-bridge cycling mechanisms in muscle contraction (Hill, 1938, Huxley, 1957). The isotonic F-V relationship provides controlled characterization of contractile properties that are widely used as parameters for musculoskeletal models, and as an applied metric to guide training in human athletes (Samozino et al., 2012; Biewener et al., 2014; Reviere et al., 2016). However, the tightly controlled conditions of the isotonic F-V relationship do not represent typical *in vivo* conditions, which are submaximal and include dynamic, time-varying varying activation, force and fascicle velocity. Yet, we do not know how *in vivo* shortening velocities relate to muscle force production during submaximal contractions.

Classic work loop techniques, pioneered by Josephson (1985), advanced our understanding of dynamic muscle contraction by measuring muscle force, fascicle strain, and work output in cyclical sinusoidal contractions (Josephson 1985). These experiments involve control of stimulation and length trajectories in *ex vivo* or *in situ* preparations and allow characterization of dynamic mechanical function of muscle (Josephson 1985; Ahn 2012). By plotting force against length over a full cycle, the area enclosed by the force-length loop corresponds to the total work output of the muscle and is therefore called a ‘work loop’. In earlier studies using the work loop technique, revealed that work loop shape changes as a function of activation phase (Tu and Dickenson 1994; Luiker and Stevens 1994; Stevens 1996; Rome and Lindstedt 1997; Askew and Marsh 1997). This foundational work also characterized the stimulation phase at which maximum muscle power output occurs (e.g., Josephson 1993, 1999; James et al., 1996; Askew et al., 1997). Other factors, like cycle frequency, strain trajectories, muscle morphology and temperature, also modulate work loop shape and therefore muscle function (Herrel et al., 2007; Asmussen et al., 1997; Luiker and Stevens 1994; Stevens 1996; Askew and Marsh 1997; Ahn 2012; Sawikci et al., 2015). These approaches provide further insight into dynamic muscle function in locomotion. Yet, classic work loop techniques use maximally activated muscles while in vivo behavior relies on a wide range of submaximal activations. As such, the isolated muscle work loop approach works well for insect muscle that often functions with a single motor unit, but less well for more complex vertebrate muscles with many more motor units. Understanding *in vivo* muscle fascicle dynamics can help inform future *in situ* and *ex vivo* experiments to allow careful determination of factors driving the link between activation, strain, and forces in locomotion.

When muscle fascicle strain and muscle-tendon force measurements are available from *in vivo* data, work loops can characterize muscle mechanical function under more dynamic *in vivo* conditions (Prilutskly et al., 1996; Roberts et al., 1997; Biewener et al., 1998a; Biewener et al., 1998b; Roberts 2001). *In vivo* work loops can characterize the complexity and variation in function of muscles in natural movements, with time-varying activation, strain, and force. These work loops can have complex shapes, due to less constrained length and activation trajectories compared to ex vivo or in situ work loops. For example, in vivo work loops in turkeys and wallabies have revealed that some distal muscles with long in-series tendons contract isometrically during force development, resulting in an “L” shaped work loop representing a ‘strut-like’ function, which enables tendinous structures to cycle elastic energy (Roberts et al., 1997; Biewener 1998b; Griffiths, 1989). *In vivo* studies have revealed that work loop shape is sensitive to the loading applied during early muscle activation, as well as the phase of activation relative to the applied load (Daley and Biewener, 2003; Daley et al., 2009; Daley 2011; Gordon et al., 2020; Schwaner et al., 2023). For example, during strenuous locomotor tasks, like incline running or obstacle navigation, guinea fowl show a more open work loop compared to steady, level walking and running, as a result of earlier force onset at longer muscle fascicle lengths (Daley and Biewener 2003; Daley 2009; Daley 2011; Gordon et al., 2020; Schwaner et al., 2023). While these *in vivo* studies provide insight into the functional range of muscle and the dynamic interactions between strain, activation, and force, they often lack insight into the underlying mechanisms because multiple factors vary simultaneously *in vivo*. Consequently, an important gap remains in the characterization of muscle dynamics through controlled benchtop experiments (*ex vivo, in situ*), and more realistic and complex conditions of *in vivo* natural movements.

The experimental conditions for the isometric F-L and isotonic F-V curves differ from *in vivo* contraction conditions because they are tightly controlled to keep length and tension constant, respectively. *In vivo* muscle fascicle length trajectories are dynamic, and force output varies depending on both activation and strain history (Josephson 1985, Edman et al., 1982; Gregor et al., 1988; Herzog et al., 1992). Furthermore, both the isometric F-L and isotonic F-V relationships are typically collected under maximal stimulation, which rarely occurs *in vivo* (Strojnik 1995). Under submaximal conditions, the plateau of the F-L curve shifts to longer lengths with decreasing activation level, and the F-V curve shifts to lower forces for a given shortening velocity (de Haan 1988; Chow and Darling, 1998; Holt and Azizi, 2014; Holt et al., 2014). These differences have important implications for interpreting *in vivo* operating ranges relative to inferred maximal capacities for force, power and shortening velocity. For example, during maximum jumping tasks, cane toads and frogs shorten their muscle fascicles across the F-L plateau, operating their muscle fascicles around the identified optimal length for maximum force output and optimal velocity for maximum power output (i.e., Holt and Azizi, 2016). However, for lower jump distances, with lower activation, *in vivo* operating ranges are poorly aligned with optimal lengths and velocities predicted by the maximal isometric F-L and F-V curves (Holt and Azizi, 2016). Previous studies have focused on muscle fascicle length operating ranges for near-maximum ballistic behaviors, and fewer have analyzed muscle force and fascicle length operating ranges during submaximal cyclical tasks (Gregor et al., 1988; Rubenson et al., 2012; Bohm et al., 2019). Therefore, it is still largely unknown how isometric F-L and isotonic F-V muscle properties relate to *in vivo* muscle fascicle operating ranges during submaximal locomotion.

Here, we investigate the correspondence between muscle isometric F-L and isotonic F-V relationships and the *in vivo* force-length dynamics and operating ranges in the guinea fowl lateral gastrocnemius (LG) muscle during walking and running. We compare measured LG isometric F-L and isotonic F-V characteristics across individuals, and directly relate each individual’s *in vivo* work loops during walking and running to their respective isometric F-L and isotonic F-V characteristics. Long-standing arguments suggest that muscle fascicles operate near optimal conditions for force and power during *in vivo* tasks (Rome 1998; Burkholder and Lieber 2001; Lieber and Ward 2011). Additionally, previous work has shown that the LG acts as a strut at high forces and as a motor at low forces during stance (Daley and Biewener 2003; Daley and Biewner 2011; Gordon et al. 2020; Schwaner et al. 2023). Therefore, we expect it to operate near optimal fascicle lengths near high forces, and near V_opt_ during rapid shortening at low forces. Our hypothesis is supported if the LG muscle fascicles operates near optimal lengths of the isometric F-L curve during *in vivo* force production, shortens across the optimal length plateau, and shortens near the optimal shortening velocity for power output according to measured isotonic F-V relationship.

## METHODS

### Animals

We used eleven adult guinea fowl (*Numida meleagris*) for this study. We divided animals into two cohorts: *in vivo* (n = 6) and non-*in vivo* (n = 5) birds. *In vivo* birds were habituated to handling and trained on the treadmill before treadmill data collection sessions. *In vivo* birds were used for *in situ* experiments following completion of the *in vivo* data collection while walking and running on the treadmill. Non-*in vivo* birds were solely used in *in situ* experiments. All procedures were licensed and approved by the University of California Institutional Animal Care and Use Committee (IACUC) (AUP 20-048). Animals were euthanized at the end of the experiments by injections of Euthasol (pentobarbital sodium) while under deep anesthesia (5%) delivered by mask.

### In vivo transducer implantation

Transducers were implanted into the lateral gastrocnemius (LG) muscle and its associated distal tendon while the animals were under surgical anesthesia (isoflurane 1.5 – 3%, mask delivery), following procedures similar to Daley and Biewener (2003, 2011). Feathers were removed from the left leg, and the surgical field was cleaned 3 times with antiseptic solution (Povidone Iodine) and alcohol wipes. Transducer leads were tunneled under the skin from the ∼20 mm incision over the synsacrum to a ∼50 mm incision over the lateral shank. Sonomicrometry crystals (1.0 mm; Sonometrics Inc, London, Canada) were implanted within the LG muscle belly in the middle 1/3^rd^ of the muscle, aligned along muscle fascicles and sutured (6-0 silk) into place. The alignment of crystals was checked post-mortem to ensure that fascicle lengths were measured accurately. Transducer signals from a sonometrics DS3 digital sonomicrometer (Sonometrics Inc, London, Canada) were checked via SonoLab software (Sonometrics Inc, London, Canada). Transducers were secured and the muscle was closed using 4-0 silk suture (Silk, Ethicon, Somerville, NJ, USA). Two bipolar electromyography (EMG) electrodes (AS 632 Teflon coated stainless steel wire: Cooner Wire Co., California, USA) were implanted in the middle third of the muscle belly. A custom-designed “E”-type stainless steel tendon buckle force transducer was placed around the common gastrocnemius tendon and secured with 5-0 silk ties. EMG and buckle transducers were connected through a 9-way micro-connector plug (580-M09, NorComp Inc., Charlotte, NC). Sonomicrometry leads were connected through an 18-way connector (ST60-18P(30), Hirose Electric Co Ltd, Downers Grove, IL, USA). Both connectors were secured using 4-0 silk suture to the skin on the bird’s dorsal synsacrum. Skin was closed using 4-0 silk suture (Silk, Ethicon, Somerville, NJ, USA). At the end of the surgery, joint centers of rotation were marked using a combination of permanent markers (black medium sharpie, Newell-Rubbermaid Office, Oak Brook, IL) and non-toxic paint (Painters, Elmer’s Products Inc, Westerville, OH).

### In vivo data collection

As part of a larger study, we recorded steady-state locomotion trials that were each 30 seconds in duration, for a total of 4 – 12 trials per individual over 1 – 3 days of experiments (2 – 6 trials per day) (Schwaner et al., 2022). Birds walked and ran on a motorized treadmill (Woodway, Waukesha, WI) at constant speeds t of 0.67 ms^−1^ and 1.56 ms^−1^, respectively. We recorded *in vivo* EMG, muscle fascicle length changes, and muscle tendon force at 5000 Hz using the USB-6363-X model National Instruments Data Acquisition board (National Instruments) through the MATLAB data acquisition toolbox (Analog Input Recorder Application, MATLAB 2021a, Mathworks, Woburn, MA). We recorded video from a sagittal plane using a high-speed camera (collected at 200 Hz, Photron VRI Veo, San Diego, CA, USA). For each bird, we selected one trial per gait condition (i.e., walking and running) for further analysis. We selected the trial in which the bird maintained steady speed for most of the trial and we had a complete dataset for the entire trial, including video, muscle force, muscle fascicle length, and EMG activity data.

### In situ data collection

For all individuals (n=11), we characterized the force-length (F-L) and force-velocity (F-V) relationships *in situ.* Birds were maintained under deep surgical anesthesia throughout the experiment (induction at 2.5-3% isoflurane, maintenance at 1.5 – 2%, mask delivery). A ∼25 mm incision was made posterior to the thigh muscles to access the sciatic nerve which was isolated and instrumented with a custom-built nerve cuff (after Naples et al., 1988) and then immersed in mineral oil. For non-*in vivo* birds, the left LG muscle was instrumented with sonomicrometry transducers (1.0 mm, Sonometrics Inc, London, Canada), positioned mid-belly along the fascicle, in the same position as for the *in vivo* birds.

For the *in vivo* birds we used the sonomicrometry crystals already placed in the previous surgery. The distal free tendon of the gastrocnemius muscle was connected to an ergometer (310C-LR, Aurora Scientific Inc., Ontario, Canada) using a custom clamp allowing measurements of muscle force and changes in muscle length. On average 21.2 mm free tendon (± 6.8 mm SD) was attached to the muscle during experiments (Table 1). The clamp was placed immediately proximal to the extra cartilaginous tissue that occurs in the tendon at the crossing of the ankle joint, while avoiding the force buckle. The medial gastrocnemius was separated from the LG by making an incision along the aponeurosis connective tissue between the two muscles.

**Table 1.** Morphological measurement. For each measurement we present the average ± 1 standard deviation (SD), range (minimum – maximum), and the number of animals these measurements were taken from.

| | Average ± SD | Range | *n* |
| --- | --- | --- | --- |
| Animal mass (kg) | 1.86 ± 0.14 | 1.68 - 2.12 | 6 |
| LG mass (g) | 9.7 ± 1.1 | 8.1 - 11.3 | 11 |
| MG mass (g) | 14.0 ± 1.3 | 12.0 - 16.3 | 11 |
| LG PCSA (mm<sup>2</sup>) | 475 ± 61 | 370 - 584 | 11 |
| Gastroc PCSA (mm<sup>2</sup>) | 1163 ± 106 | 985 - 1346 | 11 |
| Pennation Angle (°) | 22.0 ± 4.0 | 15 - 25 | 11 |
| Fascicle length, L<sub>f</sub> (mm) | 22.3 ± 1.0 | 23.5 - 27.1 | 11 |
| LG length (mm) | 98.8 ± 7.3 | 89.4 - 114.0 | 11 |
| LG thickness (mm) | 10.1 ± 1.4 | 8.3 - 12.3 | 11 |
| Free tendon length (mm) | 21.2 ± 6.8 | 9.7 - 29.8 | 11 |

For *in vivo* birds, we calibrated the E-shaped tendon force buckles before *in situ* data collection by using the ergometer to induce sinusoidal length changes on the muscle while measuring length and force. We completed the calibration of the tendon buckle before starting the F-L and F-V experiments to allow the buckle to be removed for the *in situ* experiments. We calculated a buckle calibration using a linear regression fit to the loading phase of the sinusoidal oscillations, to convert the measured strain voltage into a force in Newtons. The buckle calibrations were found to be linear within the recorded range of forces (max 38 N ± 19 SD), with a mean R^2^ of 0.95 (± 0.06 SD) and range of R^2^ between 0.85 – 0.99.

To induce muscle contractions, the sciatic nerve was stimulated using 0.2ms square wave pulses at 100 Hz (4100 High Power Stimulator, A-M Systems, Carlsborg, WA, USA). We used twitch contractions at gradually increasing voltages to determine the optimal stimulation voltage (6.5-8.5V). Stimulation trains with a 200 ms duration were used to construct the force-length relationship. Muscle force, and muscle fascicle length were recorded at 1000 Hz using a National Instruments AD board (NI USB-6363) and Igor Pro 6.4 (Wavemetrics Inc., Lake Oswega, OR, USA). To construct the F-L curve, the LG was measured under isometric conditions at a series of lengths. At each length, we first measured the passive force and length of the resting muscle, then the muscle was stimulated maximally for 200 ms (ensuring a near maximum tetanic contraction), and the maximum active isometric force and corresponding fiber length were measured for each contraction. We used 200ms stimulation to minimize muscle fatigue. Maximum tetanic force, at L_0_, was confirmed during a 400 ms contraction and was found to deviate minimally (± 4% SD) from that measured for 200 ms stimulation. The F-V relationship was determined through a series of 400 ms after-loaded isotonic contractions. Force was allowed to rise to a predefined level between 0.1 and 0.9 of F_max_ and the muscle allowed to shorten while maintaining constant force.

### Morphological Measures

Respective muscle morphological measurements can be found in **Table 1**. We obtained muscle fascicle lengths (L_fiber_), muscle length, and thickness, using calipers while the muscle was under passive tension that corresponds to an active muscle fascicle length of L_0_. These measurements were used to calculate the fractional length correction factor (L_fiber_/L_crystals_) used to convert sonomicrometry distance to the total fiber length, assuming homogeneous fiber strain. This assumption is unlikely to lead to substantial errors because the LG fibers are relatively short and L_crystals_ was large relative to L_fiber_. Crystal alignment relative to the fascicle axis was verified post-mortem, and found to be within ± 13°, indicating that errors due to crystal misalignment were <3%. The fractional length correction factor was also applied to in vivo data to calculate total fiber length and strain. We calculated the physiological cross-sectional area (PCSA) from the animal’s LG and combined LG and MG (Gastroc) muscle mass divided by muscle density times fiber lengths (L_0_) (Alexander and Vernon 1975).

Following obtaining the F-L and F-V curve for each bird, we normalized muscle fascicle length, velocity, and muscle-tendon force to the optima based on the contractile parameters obtained for that individual. Specifically, we normalized fascicle lengths by dividing instantaneous lengths by L_0_. We calculated velocity normalized to optimal fascicle length, in units of L_0_/s. The LG maximum isometric stress was calculated by dividing LG maximum isometric force (F_max_) by LG PCSA. *In vivo* stress was calculated by dividing *in vivo* measured gastrocnemius force by the combined gastrocnemius (LG and MG) PCSA, based on the simplifying assumption of uniform stress and strain between LG and MG. An estimate of total Gastroc F_max_ was calculated based on the assumption of uniform stress between LG and MG, resulting in force contributions proportional to PCSA. The total Gastroc PCSA was calculated to be 2.45X LG PCSA, therefore Gastroc F_max_ was calculated by multiplying LG F_max_ by 2.45. *In vivo* fractional force was then calculated by dividing instantaneous *in vivo* force by Gastroc F_max_ (Table 1).

### In vivo data analysis

The joint centers of rotation and several body points were digitized using DeepLabCut (version 2.2.1) (Mathis et al., 2018; Nath et al., 2019) (see Schwaner et al., 2022). Filtered marker locations (low pass 2nd order Butterworth filter, cut-off frequency of 30 Hz) were used to determine foot contact and take-off times and ankle joint kinematics. We tracked stance, swing, and stride periods based on foot contact and take-off times using the kinematics of the instrumented left leg, after Schwaner et al., 2022. Using the ankle joint kinematics, we cut strides from mid swing to mid swing. Unsteady strides resulting from accelerations, decelerations, or unexpected animal movement were excluded from analysis. Steady strides were determined based on foot velocity during stance being within 3 SD of the treadmill belt speed. We further analyzed in vivo measures of muscle force, fascicle length, activation, and kinematics in a custom written MATLAB script (MATLAB, version 2021a, MathWorks, Woburn, MA). We obtained myoelectric intensities from raw EMG signals in time and frequency using wavelet decomposition, using wavelets optimized for muscle (after Von Tscharner, 2000; Wakeling et al., 2002; Daley et al., 2009; Gordon et al., 2015; Schwaner et al., 2023). We calculated the activation threshold as +3 SD above baseline muscle activation. From these muscle intensities we obtained timing of muscle activation (EMG_on_, EMG_off_) for further analysis. To evaluate how *in vivo* fascicle length operating ranges relate to the F-L relationship, we identified five important time point during the stride cycle for further analysis: 1) the time of EMG onset T_act_, 2) time of foot on (T_on_), 3) time of 50% force rise (T _rise_), 4) time of peak force (T_peak_), and 5) time of 50% force decay (T_fall_).

### In situ data analysis

For each individual, we used an average of 12 ± 4 SD individual contractions to obtain the F-L curve. This was determined based on the shape of the force maxima of a series of contractions: we aimed to obtain contractions that would capture the parabolic shape of the F-L curve. We fitted a second-degree polynomial curve (polynomial curve fitting tool, Matlab 2022a, Mathworks, Woburn, MA) and determined the peak isometric force (F_max_) and optimum length (L_0_) from the fitted curve. To ascertain the activation dependence of L_0_ (i.e., Holt and Azizi, 2014), in one individual we measured the force-length relationship for twitch and brief 50 ms tetanic contractions, resulting in submaximal forces between a twitch and maximally tetanic contraction. We determined the corresponding peak isometric force and optimum length under these submaximal conditions. Data were then normalized to maximally activated peak force (F_max_) and optimum fiber length (L_0_) (after Holt and Azizi, 2016).

We used a series of 8 – 10 isotonic F-V contractions to determine the force-velocity relationship. We fit a standard hyperbolic equation in MATLAB to determine the F-V curve, and normalized velocity to L_0_ per second (L_0_s^−1^) to allow comparison of the F-V relationship between birds. We determined the optimal power and related optimal shortening velocity V_opt_ from the power curve (i.e, product of velocity and force). We normalized muscle shortening velocities to L_0_ (L_0_s^−1^).

### Statistical analysis

Summary values in the text are provided as mean ± SD (range: min-max), unless otherwise specified.

We used three different statistical approaches for the analysis. First, we used a paired t-test (ttest, MATLAB 2021a, Mathworks, Woburn, MA) to test the following: 1) if muscle fascicle length at EMG_on_, foot on (T_on_), and peak force (F_pk_) differed significantly from L_0_, 2) if muscle fascicle shortening velocity at T_on_ differed significantly from optimal shortening velocity (V_opt_), and 3) if muscle fascicle shortening velocity at F_pk_ differed significantly from 0. A paired t-test is appropriate because all comparisons were made within the same individual muscle and bird.

Second, we used a linear mixed-effects model (fitlme, MATLAB 2021a, Mathworks, Woburn, MA) to test relationships between the *in vivo* observed passive force and corresponding muscle fascicle strain and velocity during passive force rise. In this statistical analysis, the peak magnitude of the *in vivo* passive force during early swing was the response variable (Y). We investigated fascicle strain, fascicle velocity and gait (walk, run) as predictors, with individual as a random effect. We compared multiple models: Model 1 included only the intercept and individual as a random effect [Y ∼ 1 + (1|individual)’], which we used as a reference model and null hypothesis. This reference model was compared to 5 alternative models:

Model 2: [Y ∼ 1 + Strain + (1|individual)]

Model 3: [Y ∼ 1 + Velocity + (1|individual)]

Model 4: [Y ∼ 1 + Gait + (1|individual)]

Model 5: [Y ∼ 1 + Strain * Velocity + (1|individual)]

Model 6: [Y ∼ 1 + Gait + Strain * Velocity + (1|individual)]

We compared candidate models based on their AIC, total adjusted R2, and log-likelihood ratio tests, which supported the selection of Model 6 across all variables tested.

Finally, we used a linear mixed-effects ANOVA to test if the magnitude of the late-swing *in vivo* passive force was a significant predictor of total active force of during stance. In this test, the maximum *in vivo* active force during stance was a response variable (Z). We compared two models: Model 1 included only the intercept and individual as a random effect [Z ∼ 1 + (1|individual)’], which we used as a reference model and null hypothesis. This reference model was compared to the alternative model in which the peak late-swing passive force is a continuous linear predictor [Z ∼ 1 + Passive Force Magnitude + (1|individual)]. Based on a similar selection as described above, Model 2 was statistically supported.

## RESULTS

For each guinea fowl (n = 11), a series of isometric contractions was used to construct the passive and active curves of the LG F-L relationship (Figure 1, A-C). The maximum forces measured were 129 ± 29 N (mean ± SD), with a range of 101 – 194 Newton (N). Stress was calculated by dividing force by physiological cross-sectional area (PCSA, **Table 1**), resulting in measures of maximum muscle stress (F_max_), with a mean of 278.9± 47.2 kPa (range: 224.9 – 386.6 kPa) (**Table 2**). To compare the shape of the active and passive F-L curves across birds, we normalized the force to the maximum force (LG F_max_) and the fascicle length was normalized to L_0_ (L/L_0_) (Figure 1, D), using the morphological measures **Table 1**.

**Figure 1.**
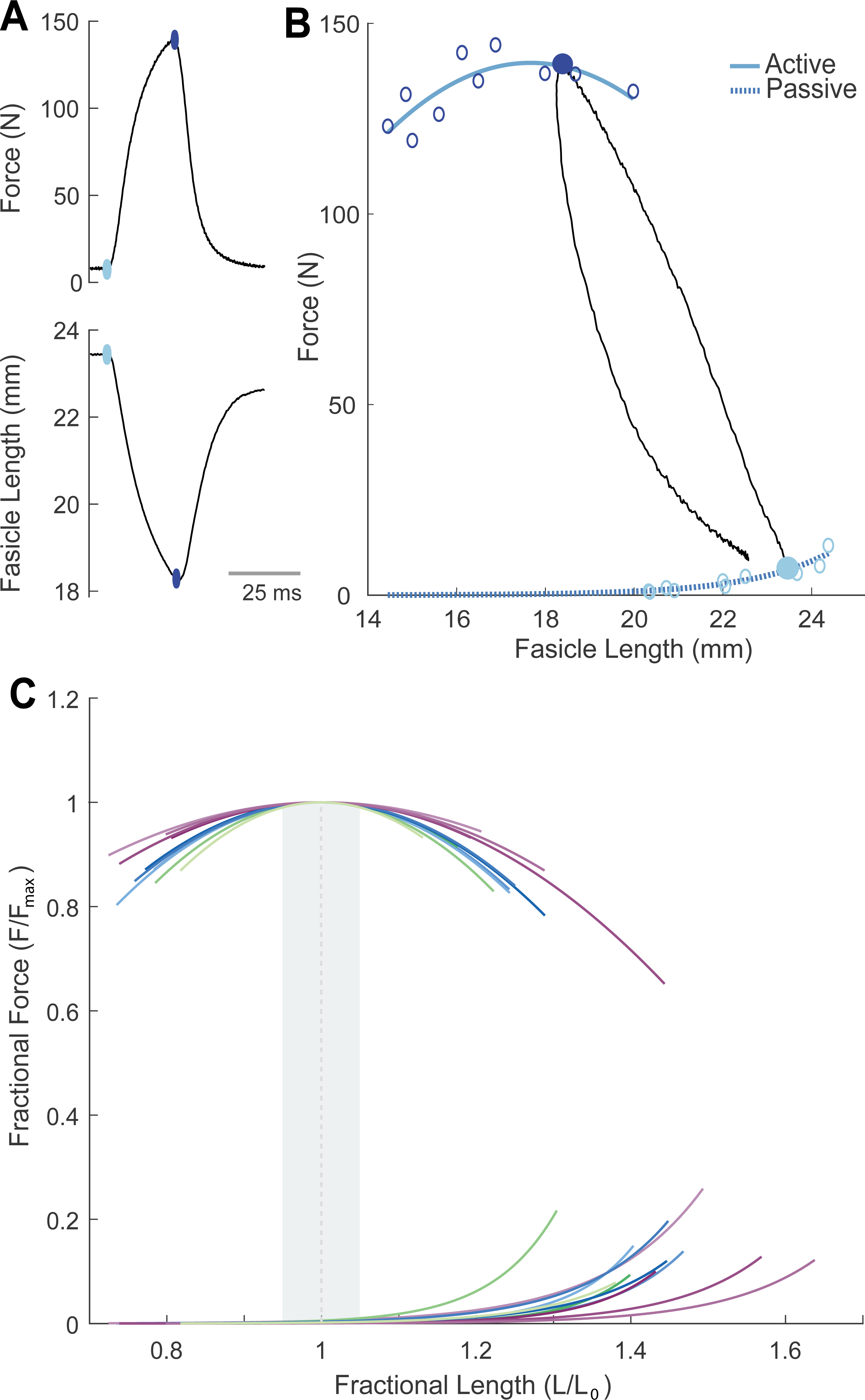
Measuring the force-length relationship. The force-length (F-L) relationship was determined through a series of 200 ms tetanic contractions at varying isometric fascicle length conditions (see methods). For each contraction (example in A), we measured the total force, and the active force (dark dot) was estimated based on the muscle fascicle shortening and fitted passive force-length relationship (light dot). **B**) After a series of contractions, separate curves were fit to the active and passive F-L data (solid and dashed lines, respectively. See methods for details. An individual contraction cycle from A is overlaid onto the F-L curve in B for illustration. **C)** We normalized data, with force as a fraction of F_max_, and length divided by L_0_ to calculate fractional length. Different colored lines correspond to the normalized F-L relationships across birds. Shaded area indicates the force plateau, defined as the region between ±5% of F_max_.

**Table 2.** Summary of key measurements from the F-L and F-V relationships across individuals. For each measurement we present the average ± 1 standard deviation (SD), range (minimum – maximum), and the number of animals these measurements represent.

| | Average ± SD | Range | *n* |
| --- | --- | --- | --- |
| L<sub>0</sub> (mm) | 19.7 ± 1.4 | 17.1- 22.1 | 11 |
| Peak force (N) | 135.97 ± 24.38 | 101.39 - 193.57 | 11 |
| Peak stress (kPa) | 278.9 ± 47.2 | 224.9 - 386.6 | 11 |
| V<sub>opt</sub> (L<sub>0</sub>s<sup>-1</sup>) | 2.9 ± 0.4 | 2.2 - 3.6 | 11 |
| P<sub>max</sub> (W kg<sup>-1</sup> muscle) | 316.4 ± 92.0 | 137.1 - 467.0 | 11 |
| V<sub>max</sub> (L<sub>0</sub>s<sup>-1</sup>) | 7.0 ± 1.1 | 5.6 - 8.6 | 11 |

\* Peak stress was determined based on peak isometric force under maximally activated conditions, fascicle length, and muscle mass

The LG Force-Velocity (F-V) relationship was constructed based on a series of individual isotonic contractions at different levels of fractional force. As an example, we show measurements from two trials from an individual bird, with force set at 0.3 and 0.9 F_max_ (Fig. 2) and illustrate where these individual measurements fall on the resulting F-V curve (Fig. 2, A-C). We multiplied force times velocity to obtain the power curve and obtained mean optimal shortening velocity (V_opt_) based on the velocity at peak power output. The mean normalized V_opt_ for all birds occurred at 2.9 ± 0.4 L_0_s^−1^ (range: 2.2 – 3.6 L_0_s^−1^) (Fig. 3). The velocity relative to V_max_ equals 0.37 ± 0.05 V_max_ (range: 0.3 – 0.5 V_max_). The mean force at V_opt_ was 0.17 ± 0.01 F_max_ (range: 0.14 – 0.18 F_max_). Summary values from the force-length and force-velocity experiments are provided in **Table 2**.

**Figure 2.**
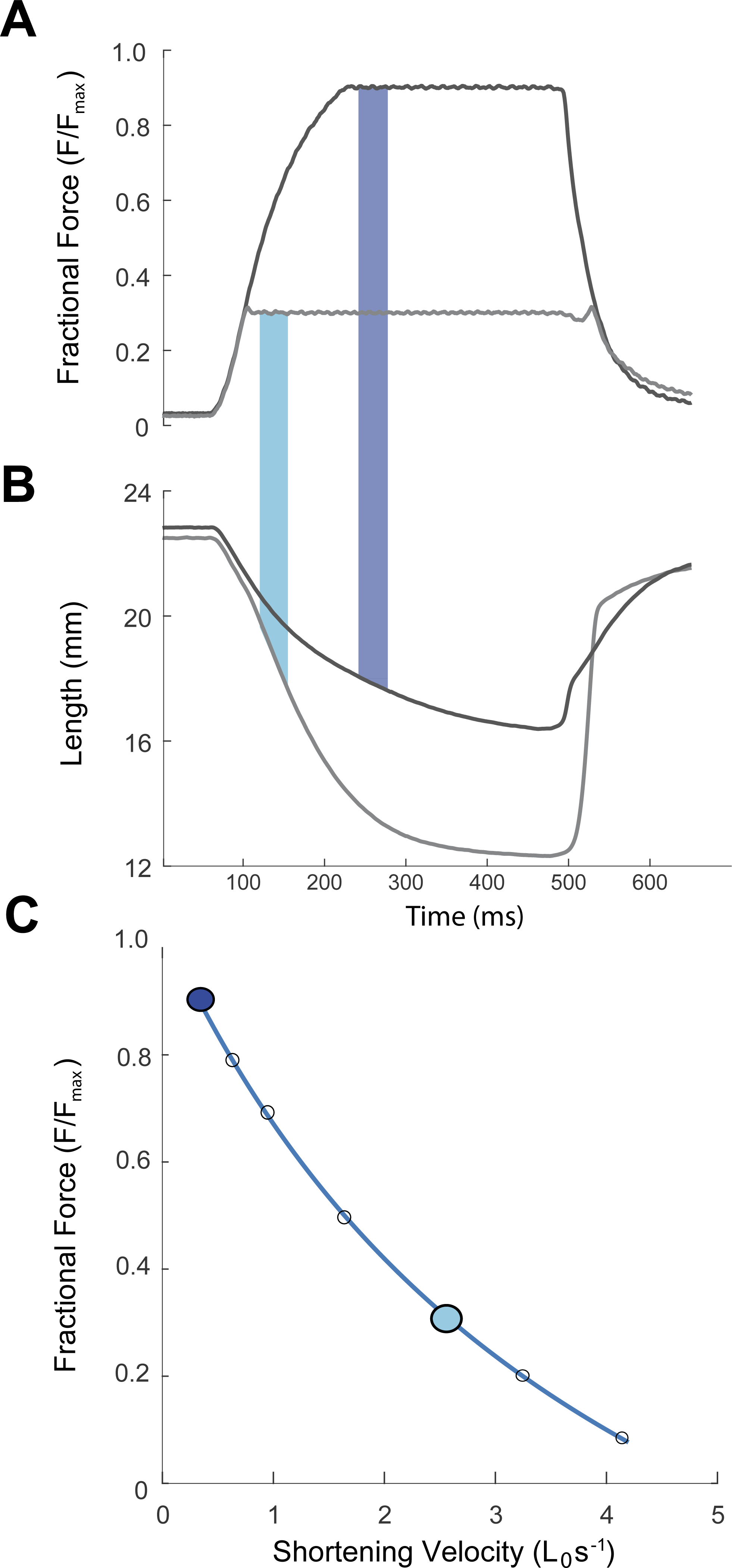
Measuring the force-velocity relationship. The force-velocity relationship (F-V) was obtained from a series of isotonic contractions at varying levels of fractional force. Two examples are shown here, for 0.3 (light) and 0.9 F_max_ contractions (A-B). The F-V relationship was fitted to data from each individual bird, using a hyperbolic curve fitting (representative shown in panel C).

**Figure 3.**
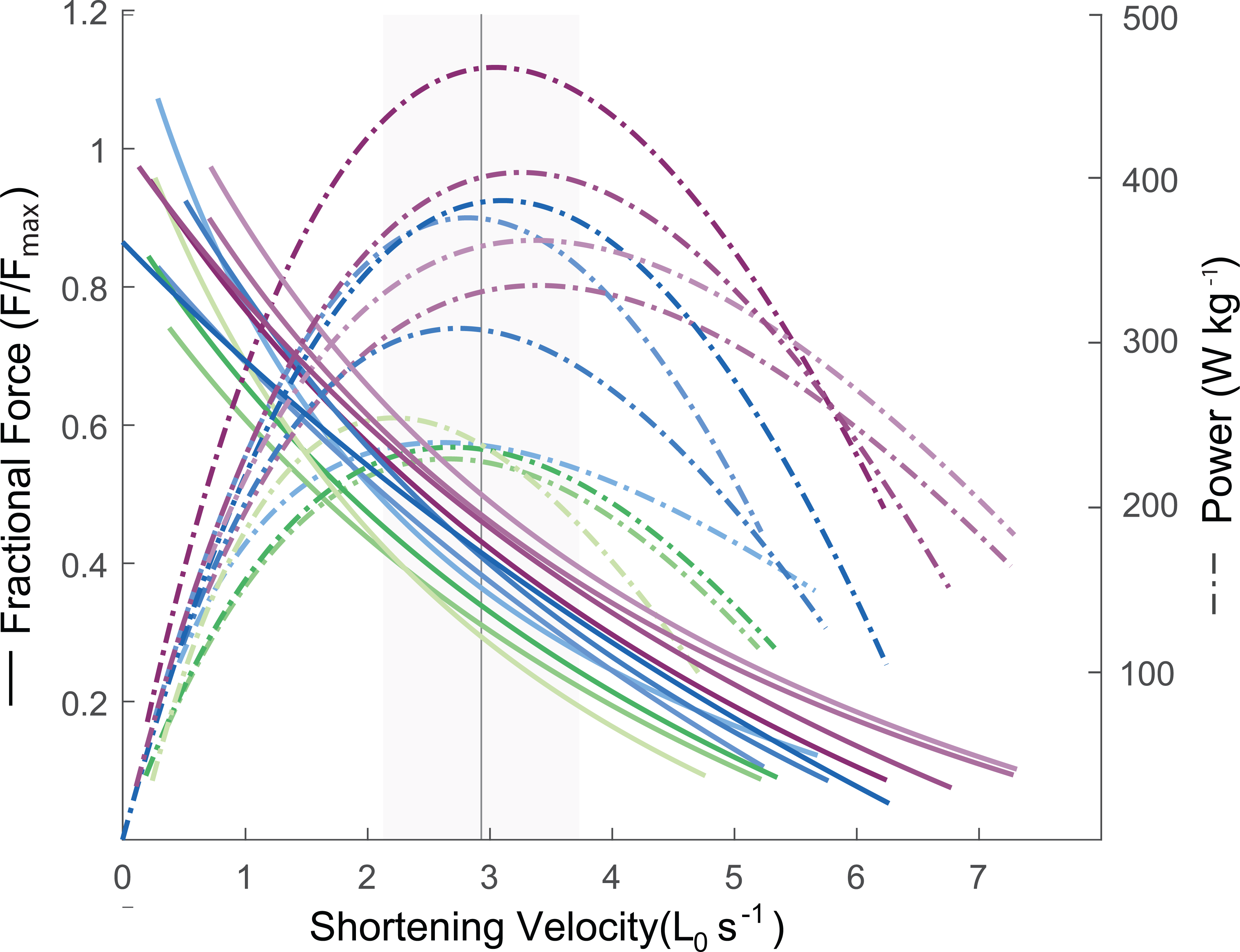
F-V relationships across individuals. The F-V relationships across individuals (n = 11) in normalized units. Power curves are shown in dashed lines in Watts per kilogram muscle. The grey vertical line indicates the mean V_opt_ (2.9 L_0_s^−1^) and the grey shading indicates the mean ± 5% V_opt_.

*In vivo* muscle force, fascicle length, and activation were used to construct muscle work loops for walking and running. A representative sequence of four strides shows typical force, length, and activation patterns over time (Fig. 4, A – B). Work loop shape is relatively repeatable across strides in steady walking and running, but does vary among individuals, as shown for two representative individuals in Figure 5. These examples represent the range of variation in work-loop shape across individuals in the current dataset. Some individuals exhibit near isometric contraction, with a narrow force peak (narrow, 5 A-B) and others exhibit substantial shortening throughout force development, resulting in an open work loop shape and higher work output (open, 5C-D). In running, total work output is higher due to a combination of higher peak force and greater shortening during force development compared to walking (Fig. 5).

**Figure 4.**
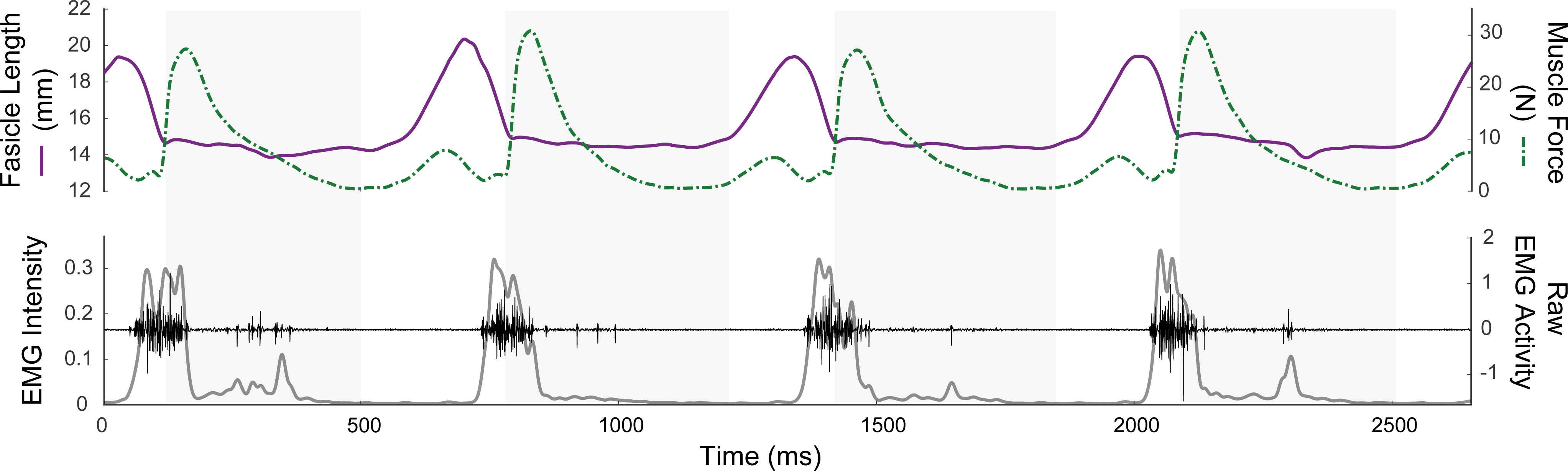
Representative *in vivo* data recording. Muscle fascicle length (top, left axis), muscle force (dashed line, top, right-hand y-axis), EMG intensity (see methods) (bottom, left axis), and raw EMG signals (bottom, right hand y-axis) for four consecutive strides for a single bird during walking. Grey shaded blocks indicate times of foot-ground contact.

**Figure 5.**
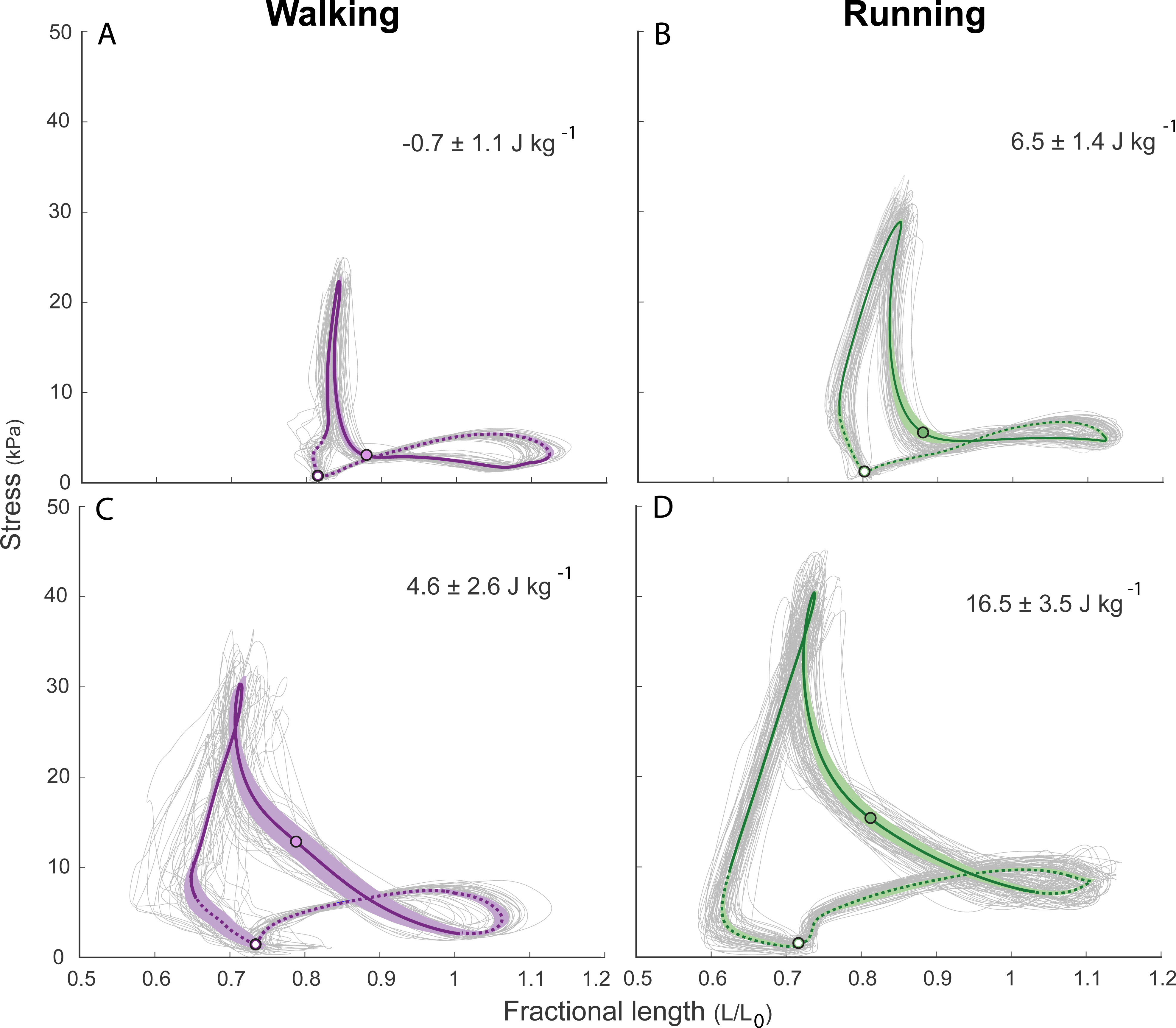
Representative work loops during walking (purple, A and C) and running (green, B and D) of two representative individuals (ind2 and ind7) All work loops of the trial are plotted in gray. Averages are represented by the colored lines. Shaded areas around these mean curves indicate the ± 95% confidence interval. Time of EMG activity is represented by the solid lines, whereas dashed lines indicate absence of EMG activity. Closed circles indicate the time of foot contact (T_on_) and open circles indicate the time of foot off (T_off_). The individuals selected span the range from the most (A) and least (B) isometric contracting LG in walking and running. Values indicate the mass specific work output (Jkg^−1^, mean ± SD).

Muscles exhibited a passive force rise during late swing during both gaits, indicated by an increase in force in the absence of EMG activity (Fig. 4 & 5). This passive force rise occurred during rapid passive stretch (Fig. 4 and 5), but at fascicle lengths near 1.0 L_0_ where no significant passive force is observed for the isometric *in situ* passive force-length curve (Fig. 1). The peak magnitude of the late-swing passive force correlates with passive muscle fascicle strain, strain rate and their interaction (strain: F-stat 40.57, p < 0.001, strain rate: F-stat 197.89, p < 0.001, strain*strain rate: F-stat 352.72, p < 0.001). Gait also has a significant effect on the peak passive force (gait: F-stat 85.94, p < 0.001). The magnitude of the passive force during swing is also a significant positive predictor of the magnitude of the active force during the subsequent stance (F-stat 125.32, p < 0.001).

To evaluate how *in vivo* fascicle length operating ranges relate to the F-L relationship, we calculated group means of normalized fascicle length and force relative to F_max_ at five time points during the stride cycle of walking and running (Figure 6, A). In both gaits, muscles are activated at lengths close to the isometric F-L plateau. In walking, length at T_act_ does not differ significantly from L_0_ (p = 0.09), whereas in running length at T_act_ is significantly longer than L_0_ (p = 0.01). Activation is followed by rapid shortening across the F-L plateau at low forces. However, significant force rise does not occur till after T_on_. At T_on_ in walking, fascicle length averaged 0.8 ± 0.1 L_0_ (range: 0.7 – 0.9), significantly shorter than L_0_ (p = 0.002). Shortly after T_on_, a sharp rise in force occurred (measured as F_rise_) leading to peak force (F_pk_). During walking, F_pk_ occurred at 0.8 ± 0.1 L_0_ (range: 0.6 - 0.9 L_0_), which is significantly shorter than L_0_ (p = 0.002). In running, fascicle length at T_on_ averaged 0.8 ± 0.1 L_0_ (range: 0.6 – 0.9 L_0_), which is significantly shorter than L_0_ (p = 0.002). At F_pk_ in running, fascicle length was 0.8 ± 0.1 L_0_ (range: 0.6 to 0.9 L_0_) which is significantly shorter than L_0_ (p = 0.002). These data indicate that *in vivo* force production during walking and running mainly occurs on the ascending limb of the isometric F-L relationships curve. Observed *in vivo* peak forces in walking and running (F_pk_) are far below F_max_. While walking F_pk_ measures 0.09 ± 0.02 F_max_ (range: 0.06 - 0.13 F_max_), whereas while running F_pk_ measured 0.13 ± 0.03 (range: 0.08 - 0.17 F_max_).

**Figure 6.**
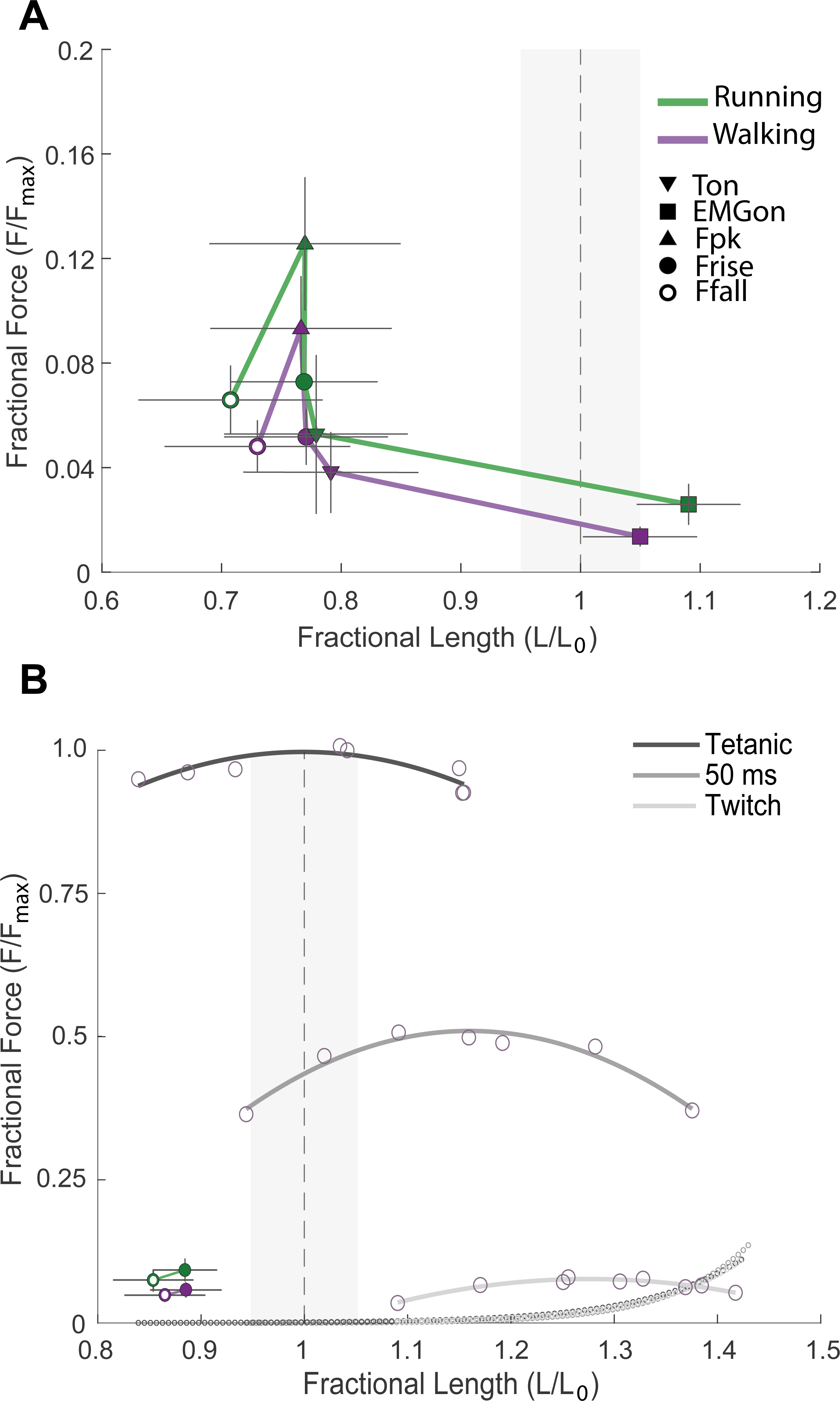
*In vivo* muscle-tendon force and fascicle strain operating ranges relative to the force-length plateau. **(A)** To summarize in vivo data, we show mean stress and strain at EMG_on_ (square), T_on_ (down pointing triangle), F_rise_ (filled circle), F_pk_ (upward triangle), and F_fall_ (open circle). Error bars indicate 95% CI. The grey shaded rectangle indicates the force plateau (±5% F_max_) of the isometric F-L relationship. The muscle is activated at lengths slightly longer than the optimum length, and rapidly shortens at low force across the plateau. No significant force rise for stance occurs until foot contact (T_on_). **B) The effect of activation on the F-L relationship.** Active F-L relationships with tetanic 200ms stimulation (dark, top), tetanic 50ms stimulation (medium, middle), and twitch stimulation (light, bottom). The corresponding passive curves are also shown but overlap. Open circles indicate individual data points from a single isometric contraction. For comparison, we also show the in vivo force and fractional length at 50% force rise and decay for walking (purple) and running (green). With decreasing stimulation and isometric force magnitude, the F-L plateau shifts rightward to longer optimal muscle lengths compared to the maximum tetanic curve.

To explore how *in vivo* velocity ranges relate to the isotonic F-V relationship, we compare *in vivo* values of velocity and force to the average F-V curve at two timepoints: at foot contact (T_on_), when force is low and power is higher, and at peak force during stance (F_pk_) (Figure 7). Muscle fascicle shortening velocity was lower at F_pk_ compared to T_on_ in both walking and running. Despite higher forces in running, shortening velocity is similar between gaits. In walking, at T_on_, shortening velocity averaged 2.3± 1.6 L_0_s^−1^ (range: 0.2 – 4.1 L_0_s^−1^), and was not significantly different from V_opt_ (p = 0.36) across individuals. At F_pk_, velocity averaged 0.02± 0.33L_0_s^−1^ (range: −0.6 – 0.6 L_0_s^−1^) which was not significantly different from zero (p = 0.89). In running, velocity at time of foot on (T_on_) averaged 2.9 ± 1.1 L_0_s^−1^ (range:1.5 – 4.3 L_0_s^−1^) and did not differ significantly from V_opt_ (p = 0.54). Velocity at F_pk_ averaged 0.03± 0.3 L_0_s^−1^ (range: 0.01 – 2.0 L_0_s^−1^) and was not significantly different from zero (p = 0.91).

**Figure 7.**
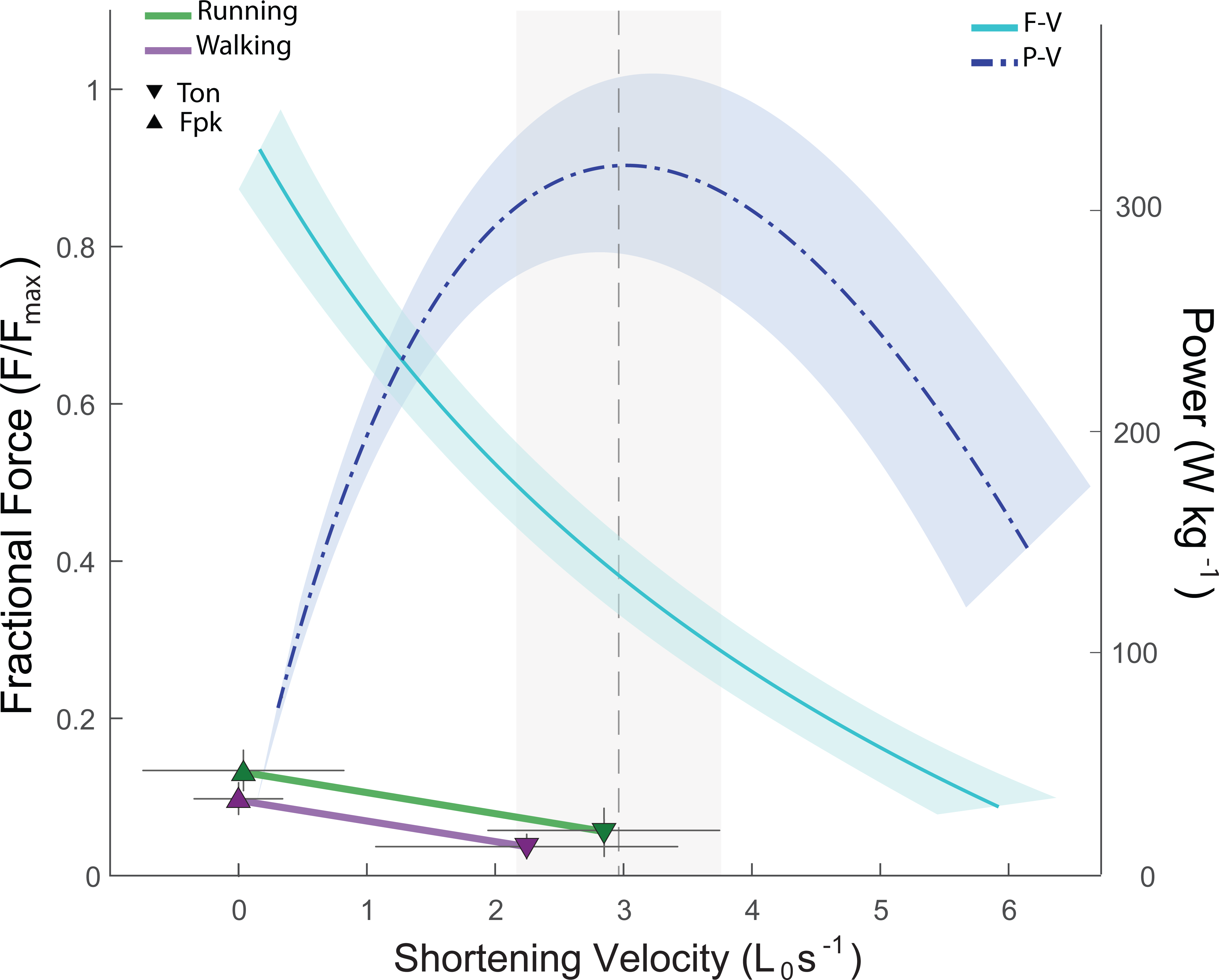
Relating in vivo operating range to the. force-velocity relationship. Mean velocity and normalized force at T_on_ (downward triangles) and F_pk_ (upward triangle) are shown along with the mean F-V and P-V relationships across individuals (light blue and dark blue, respectively). Shaded regions indicate 95% CI. The grey box and vertical dashed line indicate mean ±5% V_opt_. Muscles shorten at low velocity at F_pk_, and higher velocity at T_on_, when the muscle produces shortening work. Fractional forces are higher in running (green) compared to walking (purple).

## DISCUSSION

In the current study we compared dynamic *in vivo* operating ranges of the lateral gastrocnemius (LG) of guinea fowl to the measured *in situ* isometric F-L and isotonic F-V relationships. The muscle’s physiological properties characterized using the F-L and F-V relationship have been used to predict and interpret dynamic *in vivo* muscle dynamics (i.e., Rome 1998; Burkholder and Lieber, 2001; Lieber and Ward 2011). Based on previous work, we hypothesized that muscles operate near optimal conditions for force during early stance, and for power production in late stance during walking and running. Our findings partially support the hypothesis that muscle fascicles operate near optimal conditions— consistent with the isotonic F-V relationship, we found that LG operated at low velocity during peak force, but at velocity near V_opt_ during periods of work production in walking and running. This pattern is consistent with a previous study on turkey gastrocnemius and plantaris muscle where investigators found very low average shortening velocity (< 0.1 Vmax) at peak force (Gabaldon et. al. 2008). Although not directly reported, data from Gabaldon et al. suggest that shortening velocity is also considerably higher at T_on_ compared to peak force (see Fig. 6 in Gabaldon et. al. 2008). These findings are consistent with *in vivo* optimization for economy of forces at higher loads and power during work production, as previously suggested for the turkey gastrocnemius (Roberts et al., 1997). However, we also found that the guinea fowl LG shortens rapidly at low forces across the isometric F-L plateau and produces the highest forces at lengths around 0.8 L_0_. This indicates that the LG mainly develops force at lengths well below the optimal length predicted based on the isometric F-L plateau. We also observe that the F-L curve shifts rightward (to longer lengths) with submaximal activation, therefore the in vivo operating ranges correspond to lengths further down the ascending limb of the submaximal F-L curve (Fig. 6B).

In series tendons of pennate muscles facilitate decoupling of muscle fascicle and muscle tendon unit (MTU) dynamics (Roberts et al., 1997; Azizi and Roberts 2014; Schwaner et al., 2021). Previous work suggests that bi-articular LG MTU acts to control energy transfer across joints and provide ankle propulsion (i.e., Gregoire et al., 1984; Zajac et al., 2002; Lichtwark and Wilson, 2006; Schwaner et al., 2021). Here we find that guinea fowl LG work loops exhibit an L-shape (Fig 3.A), which suggests mostly strut-like function of the muscle fascicles during stance. At the swing-stance transition, LG muscle fascicle shortening work occurs simultaneously with energy absorption at the ankle joint (Daley and Biewener 2003; Daley and Biewener 2007; Daley et al., 2009), suggesting that muscle fascicles shorten against the tendon while the tendon stretches under load. During early stance, the ankle joint absorbs energy, therefore the positive work done by LG would not directly power joint work. It is interesting that the fascicles shorten near V_opt_ even when the muscle is not directly powering joint or limb propulsion. In late stance, positive power occurs at the ankle, yet muscle fascicles contract near isometrically, enabling elastic energy recoil in the tendon, as observed in turkeys and wallabies (Roberts et al., 1997; Griffiths, 1989). These findings suggest that in steady gait, the guinea fowl LG mainly controls energy transfer across joints, rather than directly actuating the ankle joint. Considering that the LG is mainly regulating force during stance, it is perhaps surprising that the LG muscle fascicles do not operate on the F-L plateau for optimum economic force production.

There may be a potential functional benefit to operating at lengths shorter than optimal in steady state. Obstacle perturbations sometimes require sudden increase in muscle force development at longer fascicle lengths, for example, in uneven terrain (Daley and Biewener 2011). The LG starts shortening near L_0_ at time of EMG onset, yet force production in steady gait occurs almost entirely on the ascending limb of the isometric F-L curve (0.7-0.8 L_0_). Operating at lengths on descending limb of the F-L curve is thought to be a strategy for avoiding the muscle damage that may arise from a rapid stretch due to an unexpected perturbation. Here the system needs to balance what is likely a complex trade-off between economic force generation, the ability to develop force rapidly to resist a minor stretch (which increases with length), and avoidance of instabilities associated with long sarcomere lengths. Although this pattern may not be optimal for steady treadmill locomotion, the additional force potential may provide intrinsic mechanical stability in the presence of rapid unexpected perturbations. In the presence of obstacles, guinea fowl LG experiences earlier loading due to earlier foot contact, resulting in an earlier and rapid rise in force, reaching a higher peak force (Daley and Biewener, 2011; Gordon et al., 2020; Schwaner et al., 2023). The muscle fascicles continue to shorten throughout the perturbation, therefore increased force is not directly related to fascicle stretch. However, force development occurs at longer lengths in obstacle steps. The current findings suggest some of the increase in force in obstacle steps arises from higher force potential when the fascicles operate at lengths on the F-L plateau (Fig 8). A steady state work loop trajectory that operates at shorter than optimal lengths provide potential for a rapid, intrinsic increase in muscle force when the operating length of the muscle fascicles is increased by a sudden abbreviation of the swing phase, reacting more rapidly than possible through reflexive changes in activity (Loeb et al., 1999; Daley et al., 2009). Therefore, the shortening of LG across the F-L plateau in late swing may provide a mechanical safety factor against unexpected perturbations that require rapid force development in response to varied timing of foot contact.

**Figure 8.**
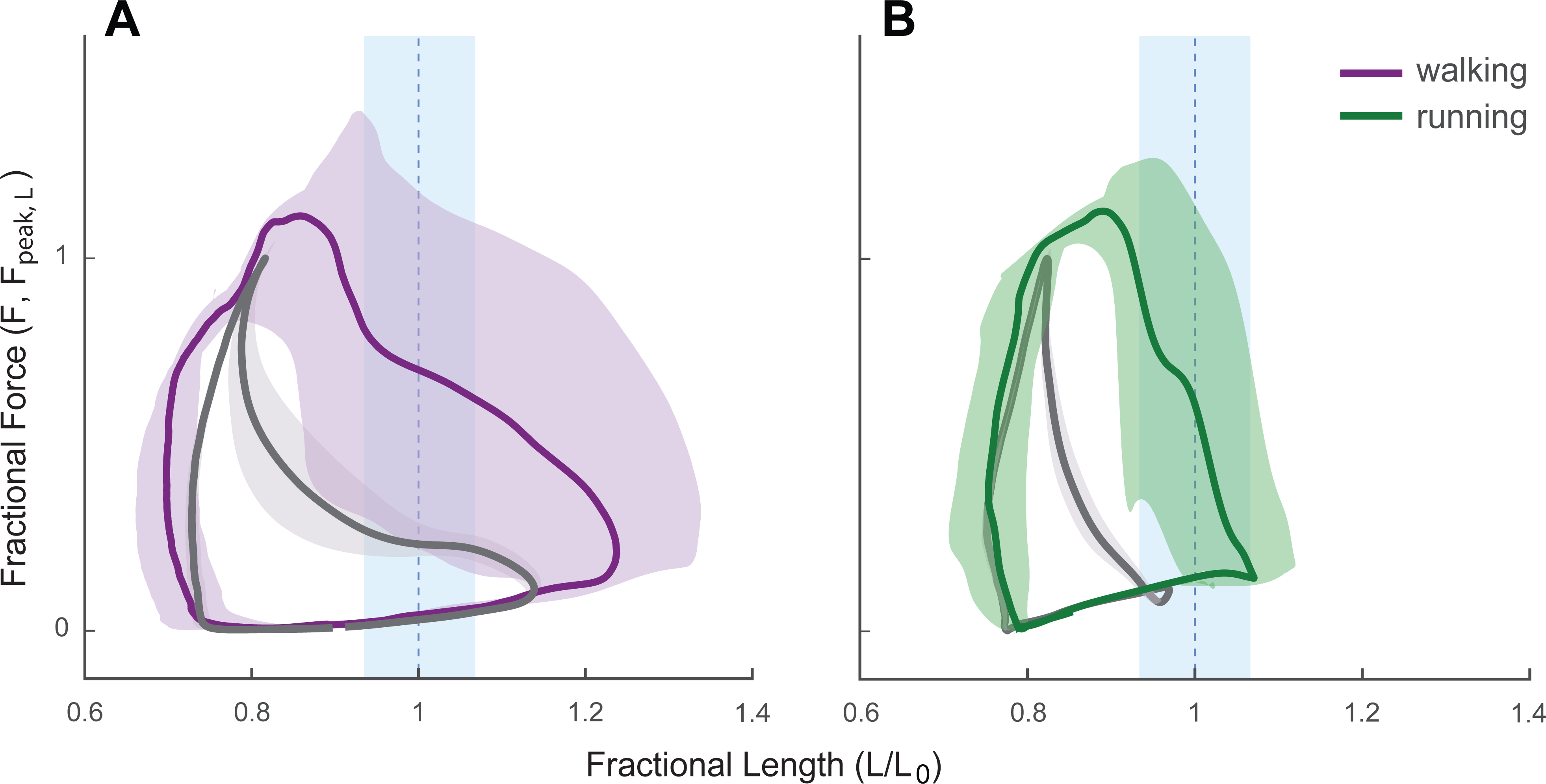
LG work loops in obstacle strides (color) compared to level strides (grey) in walking (A) and running (B) Data from published studies on walking (A, Ind 3) and running (B, Ind 6) over obstacles (Gordon et al., 2020; Schwaner et al., 2023). Force is normalized relative to peak force during steady level gait at the same speed because F_max_ is unknown. Fascicle length has been normalized such that peak force occurs at 0.8, as found in the present study. However, L_0_ was not directly measured for these individuals. Shading represents 95% CIs. Obstacles elicit earlier and steeper force rise, at longer fractional lengths compared to level. Peak forces occur on estimated F-L plateau in obstacle strides. This suggests that the low forces in late swing in steady strides create a safety buffer, with potential for the muscle to rapidly generate higher forces in response to unexpected perturbations, without a change in muscle activation.

Shortening across the F-L plateau may occur in cyclical tasks that involve abrupt transitions in external loading with uncertain timing. Shortening across the F-L plateau is also observed in frog plantaris and semimembranosus in hopping, the rat medial gastrocnemius during totting, and the sternohyoid muscle during swallowing (Ahn et al., 2018; Holt and Azizi 2016). In contrast, during jumping tasks, frog plantaris muscles primarily operate on the descending limb of the F-L curve (Azizi and Roberts, 2010). These patterns may relate both to the mechanical demands of the task and the uncertainty in externally applied loads. Ballistic movements such as jumping may involve more predictable loading patterns compared to cyclical tasks that involve abrupt load transitions in contact with the external environment.

An important difference between the dynamic *in vivo* and *in situ* experimental approaches presented here is the level of muscle activation. Maximum muscle activation is typically used to determine the F-L and F-V relationships, but is rarely observed in nature (Walmsley et al., 1978; Strojnik 1995), most likely because of energetics cost and the potential of irreversible muscle damage (Crameri et al., 2007). Additionally, there is a known dependence of L_0_ on activation, with the F-L shifting rightward, toward longer lengths, with decreasing activation (e.g., Holt and Azizi, 2014; Holt and Azizi, 2016). We found a similar shift in the F-L curve with submaximal stimulation (Fig. 6,B). This suggests that *in vivo* operating ranges occur even further down the ascending limb of the submaximal F-L plateau than suggested by the maximally stimulated F-L curve (Fig 6).

The isometric F-L and isotonic F-V relationships allow comparison of muscle contractile properties across under standard and controlled conditions, but they do not capture nonlinear interactions between strain trajectory and activation dynamics that influence force and work output during dynamic tasks (Askew and Marsh 1998; Perrault et al., 2003; McGowan et al., 2013). We found that a passive force rise (i.e., in absence of EMG activity) occurs in early swing (Figure 3 and 4) in walking and running. This passive force rise occurs at fascicle lengths that do not result in measurable muscle passive forces in situ (Figure 1 and 5). This suggests that the passive force-length relationship under static conditions cannot capture these observed forces, which might instead relate to dynamic viscoelastic properties or history dependent effects, such as short-range stiffness, stretch-induced residual force enhancement, and passive force enhancement (Edman et al., 1982; Edman 2012; Herzog 2018, Seiberl et al., 2015; Hahn and Riedel 2018). Viscoelastic properties and history dependent effects can provide stability in response to perturbations as active control can tune intrinsic properties (Gregor et al., 1988; Loeb et al., 1999; Lappin et al., 2006). We find that the magnitude of the active force in the stance phase has a positive correlation with the magnitude of the passive force during swing, suggesting the importance of contributions of dynamic strain and strain rate histories on passive and active force development, even during steady state *in vivo* conditions. Viscoelastic elements, like titin, contribute to muscle force and work during steady-state locomotion, thereby assisting in stabilizing gait (Zajac et al., 2003; Hessel et al., 2021). However, we note that the Achilles tendon is the common tendon for the LG, IG and MG in guinea fowl, and therefore contributions from IG or MG, via either direct or lateral transmission of force, could potentially contribute to the apparent passive forces. Yet, no significant MG EMG activity is observed in late swing during walking and running in guinea fowl (Gordon et al., 2015). Given these results, it is critical that the approaches to studying isolated muscles advance beyond the standard historical protocols. While direct measures of muscle force and length *in vivo* may not be practical for all systems, recent technical advances provide the potential to bridge the gaps in current understanding by using real-time feedback control to allow isolated muscles to interact dynamically with models (physical or virtual) (Robertson and Sawicki, 2015; Richards and Eberhard 2020; Mendoza et al. 2023). Further work and new approaches are needed to understand how the nonlinear interactions between activation and strain trajectory influence muscle force and work output during dynamic contractions *in vivo*.

This study provides a novel comparison of *in vivo* muscle operating forces, fascicle lengths, and velocities to optimum *in situ* characterized lengths and velocities. We show that the guinea fowl LG muscle fascicles shorten near optimal velocities during force development *in vivo*, but do not operate on the F-L plateau during force development in walking and running. We also found a rise in passive force in LG during early swing at fascicle lengths that correspond to negligible passive forces *in situ*. This finding suggests that *in vivo* force-length dynamics are substantially influenced by dynamic strain effects such as viscoelasticity, short-range stiffness, and other lengthening force enhancement. Our study provides new evidence that the static nature of the F-L properties makes intrinsic muscle properties less predictive of *in vivo* behavior than previously thought. Direct comparison of isometric F-L and isotonic F-V properties to *in vivo* muscle force-length dynamics and operating ranges also suggest the need to further examine how dynamic interactions between strain and activation influence muscle force and work output *in vivo*. Complex interactions between strain and activation are especially important during unsteady locomotor tasks, where muscle tendon force and work output may be influenced by viscoelastic and other history-dependent aspects that are not captured in isometric F-L and isotonic F-V experiments.

## ACKNOWLEDGMENTS

The authors would like to acknowledge Elizabeth Mendoza for expert advice during *in situ* data collection. The authors would like to thank Adrien Arias, Vivian Chong, Brooke Christensen, Catalina Dentzel, Kamila Karimjee, Zhiji Li, Elizabeth Mendoza, and Daisey Vega for help with *in vivo* data collection, and Chris Wagner for helping with the *in vivo* experimental set-up.

## DATA AVAILABILITY

The data underlying this research article will bemade available through DataDryad upon acceptance by the journal. Data include *in vivo* muscle fascicle length, muscle-tendon force, and activity recordings of running and walking guinea fowl, as well as their muscle physiological properties of isometric F-L and isotonic F-V. Additionally, the data underlying our statistical tests are presented in a spreadsheet. Link to data repository: [data will be made available upon acceptance by journal].

